# *Rhipicephalus microplus* and its vector-borne haemoparasites in Guinea: further species expansion in West Africa

**DOI:** 10.1101/2020.11.12.376426

**Authors:** MT Makenov, AH Toure, MG Korneev, N Sacko, AM Porshakov, SA Yakovlev, EV Radyuk, KS Zakharov, AV Shipovalov, S Boumbaly, OB Zhurenkova, YaE Grigoreva, ES Morozkin, MV Fyodorova, MY Boiro, LS Karan

## Abstract

*Rhipicephalus microplus* is an ixodid tick with a pantropical distribution that represents a serious threat to livestock. West Africa was free of this tick until 2007, when its introduction into Benin was reported. Shortly thereafter, the further invasion of this tick into West African countries was demonstrated. In this paper, we describe the first detection of *R. microplus* in Guinea and list the vector-borne haemoparasites that were detected in the invader and indigenous *Boophilus* species.

In 2018, we conducted a small-scale survey of ticks infesting cattle in three administrative regions of Guinea: N`Zerekore, Faranah, and Kankan. The tick species were identified by examining their morphological characteristics and by sequencing their COI gene and ITS-2 gene fragments. *R. microplus* was found in each studied region. In the ticks, we found DNA of *Babesia bigemina*, *Anaplasma marginale, Anaplasma platys*, and *Ehrlichia spp*. The results of this study indicate that *R. microplus* was introduced into Guinea with cows from Mali and/or Ivory Coast.

## Introduction

*Rhipicephalus microplus* (Canestrini, 1888) (formerly *Boophilus microplus*) is an ixodid tick species with a one-host life cycle that parasitizes a variety of livestock species. *R. microplus* is the most commercially important tick species distributed in tropical and subtropical regions of the world (Frisch 1999; Labruna et al. 2009). In Africa, this tick has been established in much of southern and eastern Africa, including such countries as South Africa, Mozambique, Zimbabwe, Malawi, Zambia, Tanzania, and Kenya (Walker et al. 2003). Since 2007, *R. microplus* has been detected in West Africa, specifically Ivory Coast (Madder et al. 2007) and Benin (de Clercq et al. 2012; Madder et al. 2012). Soon after these findings, several papers confirmed the further invasion of the species in West Africa: *R. microplus* was detected in Burkina Faso, Togo, Mali (Adakal et al. 2013), Nigeria (Kamani et al. 2017), and Cameroon (Silatsa et al. 2019).

The rapid spread of *R. microplus* in West Africa entails a negative impact on the livestock sector. Heavy infestation of cattle with this tick causes weight loss, reduced milk production and increased cow mortality (Corrier et al. 1979; Guerrero et al. 2007; Madder et al. 2011; Rocha et al. 2011; da Silva et al. 2013). Furthermore, *R. microplus* causes considerable losses in cattle as a vector of several pathogens, including *Babesia bigemina*, *Babesia bovis, Anaplasma marginale*, and *Ehrlichia ruminantium* (Guerrero et al. 2007; Biguezoton et al. 2016).

In the Guinea sub-genus, *Boophilus* was represented by three species: *R. annulatus* (Say, 1821), *R. decoloratus* (Koch, 1844), and *R. geigyi* (Aeschliman & Morel, 1965) (Konstantinov et al. 1990; Walker et al. 2003; Bouaro et al. 2013). *R. microplus* has not been detected in Guinea previously. In this paper, we confirm the presence of *R. microplus* in Guinea and list the vector-borne haemoparasites that were detected in the invader and indigenous *Boophilus* species.

## Methods

In April-May 2018, we conducted a small-scale survey of ticks infesting cattle in three administrative regions of Guinea: N`Zerekore, Faranah, Kankan. These regions are located on the eastern part of Guinea and adjoin countries where the presence of *R. microplus* has been confirmed previously (Mali, Ivory Coast). We collected ticks from freshly slaughtered cattle or cattle prepared for slaughtering in slaughterhouses in each region. Ticks collected from a cow were placed in a separate tube. The sampled ticks were identified to stage and species according to their morphological characteristics (Walker et al. 2003) and were pooled into groups of one to two for *Rhipicephalus* and *Hyalomma* species and from one to five for *Amblyomma variegatum* Fabricius, 1794. Pools were formed by species, sex, and animal host. Ticks were later washed with 70% alcohol and then rinsed twice with 0.15 M NaCl solution. Each pool was homogenized with TissueLyser LT (Qiagen, Hilden, Germany) in 500 μl of 0.15 M NaCl solution. Thereafter, DNA/RNA was extracted from 100 μl of the 10% tick suspension using a commercial AmpliSens RIBO-prep kit (Central Research Institute of Epidemiology, Moscow, Russia) following the manufacturer’s instructions.

To check the tick species diagnoses based on morphological characteristics, we sequenced the COI gene fragments for each *R. microplus* and for several voucher specimens for other tick species using primers described previously (Geller et al. 2013; Makenov et al. 2019). Additionally, we sequenced the ITS-2 gene fragments for *R. microplus* specimens using the following primers: 58S F3/1: 5’-GGGTCGATGAAGAACGCAGCCAGC-3’ (Fukunaga et al. 1995) and ST-ITS2-R2: 5’-AACCGAGTRCGACGCCCTACCA-3’ (self-designed).

Since the primary topic of this paper is the invasion of *R. microplus* in Guinea, we investigated the subgenus *Boophilus*; other tick species (*A. variegatum* and *Hyalomma truncatum*, Koch, 1844) were excluded from further consideration. Additionally, *R. microplus* parasitizes only cattle and wild ungulates; consequently, we focused this paper on haemoparasites of cattle and did not consider other vector-borne pathogens. We screened for the presence of *Babesia spp.* with primers Bs1 and Bs2 (18S rRNA gene) according to Rar et al. (2011) and by qPCR (Michelet et al. 2014) (see primers and probes in Table 1). Additionally, we used qPCR to screen *Theileria annulata* (Michelet et al. 2014) (see primers and probes in Table 1). To detect bacteria of the Anaplasmataceae family, we used primers Ehr1, Ehr2, Ehr3, and Ehr4 (16S rRNA gene) according to (Rar et al. 2010). The identification of *Anaplasma marginale* was additionally confirmed by sequencing of the groEL gene fragment: the assay was conducted in a nested format with primers HS1-f and HS6-r (Rar et al. 2010) in the first round and primers HS3-f HSVR in the nested reactions (Liz et al. 2000). Furthermore, we employed qPCR to screen *A. marginale, A. ovis*, and *A. centrale* (Michelet et al. 2014) (see primers and probes in Table 1).

**Table 1.**
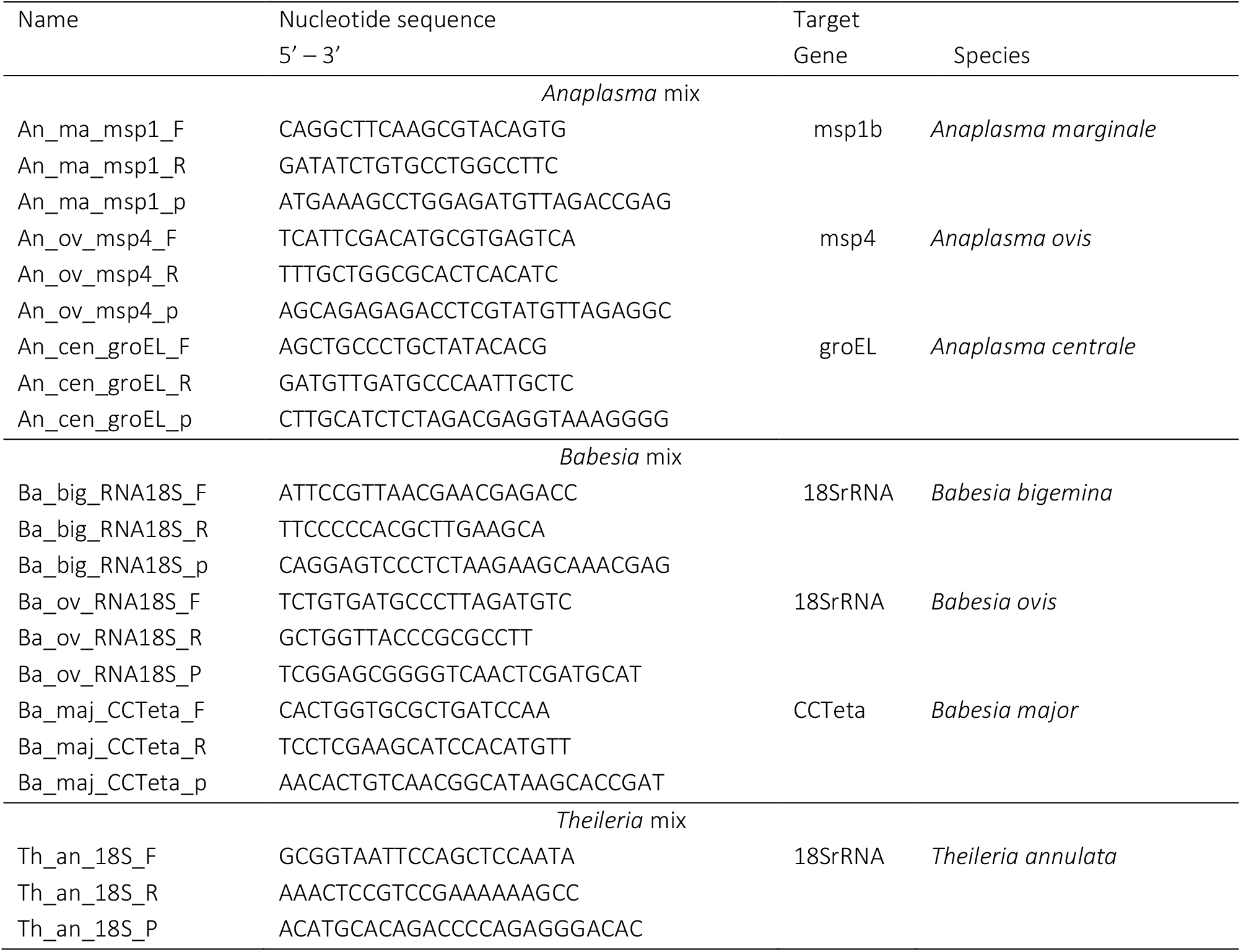
Oligonucleotide sequences of the primers and probes used in qPCR amplification of *Anaplasma spp.* and *Babesia spp*.(according to Michelet et al (Michelet et al. 2014))

Sequencing was conducted using BigDye Terminator v1.1 Cycle Sequencing kit (Thermo Fisher Scientific, Austin, TX, USA) on an Applied Biosystems 3500xL Genetic Analyzer (Applied Biosystems, Foster City, CA, USA). The obtained sequences were deposited in NCBI GenBank under the following accession numbers: MT107445-MT107461 – *R. microplus* COI gene fragments; MT107428, MT107430, MT107433-MT107444 – *R. geigyi* COI gene fragments; MT107429, MT107431-MT107432 – *R. annulatus* COI gene fragments; MT112107-MT112110 – *R. microplus* ITS-2 gene fragments; MT112111-MT112112 – *R. geigyi* ITS-2 gene fragments; and MW042697-MW042698 – *A. platys* 16S rRNA gene fragments; MW042699-MW042705 – *A. marginale* 16S rRNA gene fragments; MW042706 – *Ehrlichia sp*. 16S rRNA gene fragments; MW054555-MW054558 – *Ehrlichia sp.* groEL gene fragments; and MW049239-MW049240 – *Babesia bigemina* 18S rRNA gene fragments.

## Results

A total of 40 cows were examined: 17 in Faranah, 10 in N`Zerekore, and 13 in Kankan. In total, 561 ticks of six species were collected (Table 2). The sub-genus *Boophilus* was represented in the sample by three species, including *R. microplus*, *R. annulatus*, and *R. geigyi* (Table 2). *R. microplus* was found on cows from eight different villages located in all three studied administrative regions (Fig. 1). We found either adult ticks or eggs: one female of *R. microplus* began oviposition in the tube.

**Fig. 1.**
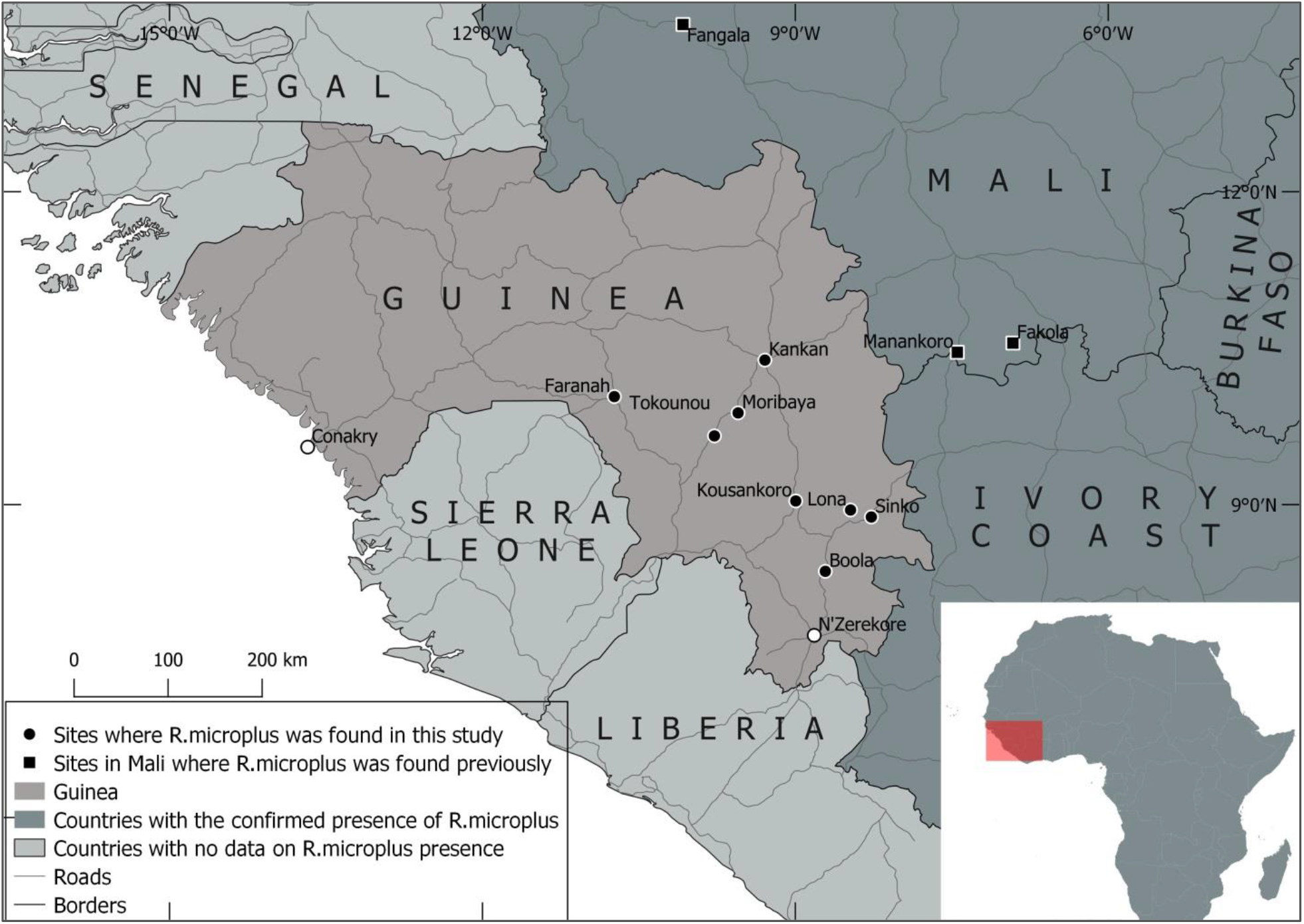
*Rhipicephalus microplus* in Guinea and surrounding countries

**Table 2.**
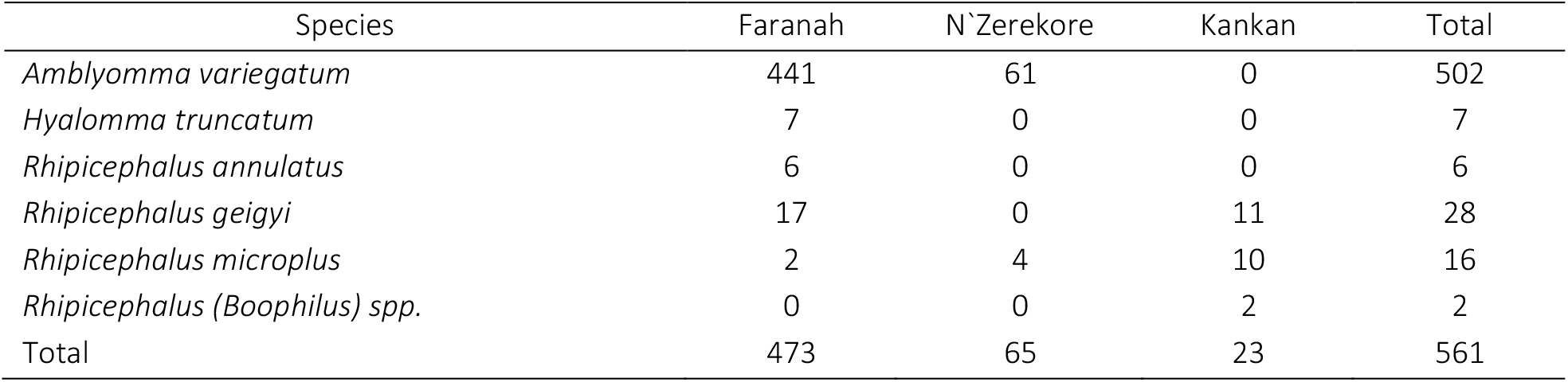
Number of ticks collected from cattle in three administrative regions of Guinea in April-May 2018

Sequencing of the COI and ITS-2 genes confirmed the morphological identification of *R. microplus*. BLAST analysis of these sequences revealed 100% identity with the sequences of *R. microplus* available in the GenBank database, including the COI gene sequence of *R. microplus* from Benin (accession number KY678120).

Screening of *Babesia spp*. showed the presence of *Babesia bigemina* DNA in *R. microplus* only (Table 3). Four infected ticks were collected from two cows in two different villages in the Kankan region. Furthermore, *B. bigemina* was also detected in samples of eggs of *R. microplus*. Two other species of protozoan haemoparasites, *Theileria annulata* and *Theileria parva* were not found in the studied ticks (Table 3).

**Table 3.**
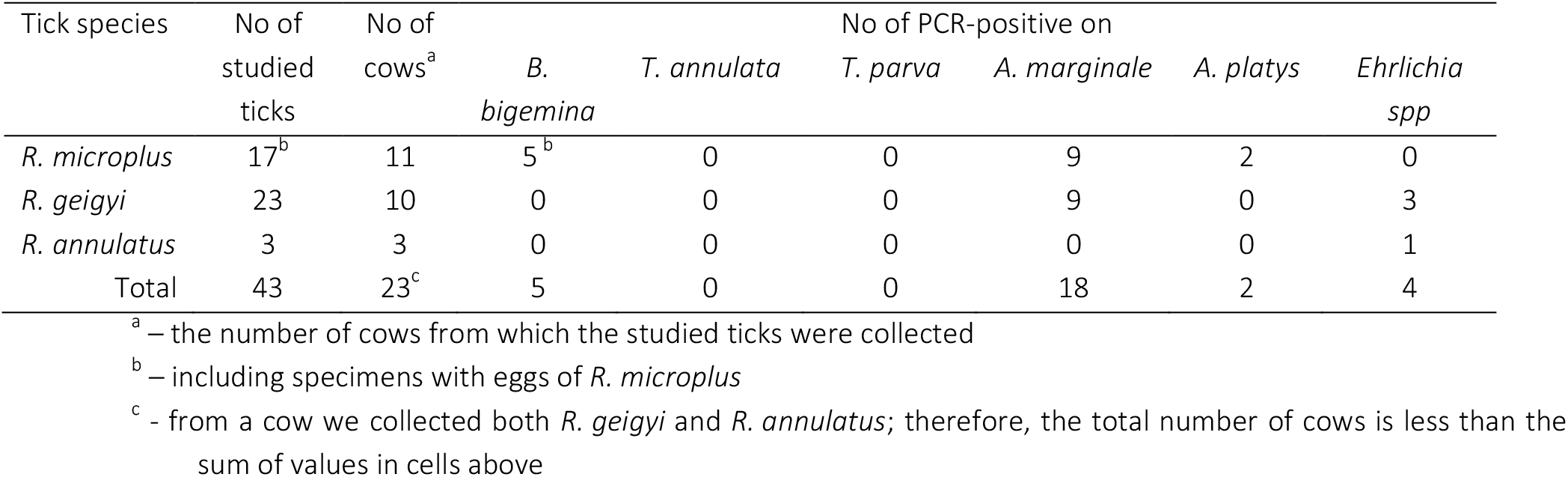
Results of PCR screening on *Babesia bigemina, Theileria annulata, Theileria parva* and bacteria of the Anaplasmatacea family in ticks of the *Boophilus* sub-genus

*Anaplasma marginale* was found in *R. microplus* and in *R. geigyi* collected in the N`Zerekore and Kankan regions. Additionally, two females of *R. microplus* (from Faranah and Kankan) were infected with *A. platys* (Table 3). We also found *Ehrlichia spp.* in four ticks collected from a host cow in Faranah. The obtained sequences of groEL and 16S rRNA genes enable us to identify the pathogen at the genus level only. Most similar sequences from GenBank belong to uncultured *Ehrlichia* isolated from ticks in China and Tajikistan: 97.7% identity on the groEL gene with *Ehrlichia spp*. isolated from ticks in Tajikistan (GenBank accession number KJ930191) and 99.8% identity on the 16S rRNA gene with *Ehrlichia spp*. isolated from ticks in Niger, Malaysia, China, and Thailand (GenBank accession numbers AF311968, KY046298, KJ410255, and AF497581, respectively).

## Discussion

The spread of *R. microplus* in West African countries is associated with the transfer of livestock between countries (Madder et al. 2012; Adakal et al. 2013). Evidently, *R. microplus* also appeared in Guinea with cattle imported from neighbouring countries. Guinean veterinarians confirmed that farmers buy cows every year in Mali, Ivory Coast, and Senegal. In particular, we detected *R. microplus* in Beyla Prefecture (N`Zerekore region), and since 2011, local farmers have purchased cattle, including cows of the breed Girolando, from Mali with transfers via the Ivory Coast. According to Adakal et al (2013), *R. microplus* was found in three villages in Mali in 2011. Two of these villages (Manankoro and Fakola) are located approximately 200-250 km from Beyla Prefecture. Furthermore, the N`Zerekore and Kankan regions share a border with the Ivory Coast (Denguele and Woroba Districts), where *R. microplus* was also detected (Boka et al. 2017). Therefore, we hypothesize that *R. microplus* was introduced into Guinea with cows from Mali and/or Ivory Coast.

Previously, it was shown that the introduction of *R. microplus* to new territories could lead to displacement of the native *Boophilus* species (Macleod and Mwanaumo 1978; Wedderburn et al. 1991; Berkvens et al. 1998; Tønnesen et al. 2004). In West Africa, the displacement of *R. decoloratus* and *R. geigyi* was observed in the Ivory Coast (Madder et al. 2011; Boka et al. 2017) and Benin (De Clercq et al. 2012). We found ticks of the subgenus *Boophilus* in samples from nine Guinean villages, and in eight of them, *R. microplus* was detected. The simultaneous presence of *R. microplus* with other species of the subgenus was observed in Tokounou village (Kankan region) and in Faranah. Due to a lack of sampling efforts, we could not assess the replacement of native species of the *Boophilus* sub-genus. However, the detection of *R. microplus* and *R. geigyi* in the same village indicates that the displacement process had not begun or was in an early stage.

De Clercq et al. (2013) calculated a climate suitability map for *R. microplus* in West Africa. According to the model of these researchers, the species’ potential range covers the southern part of Guinea: N`Zerekore region and parts of Kankan, Faranah, and Kindia regions (De Clercq et al. 2013), and our findings of *R. microplus* correlate well with the predicted suitable area. Liberia and Sierra also belong to territories with high climate suitability for *R. microplus* (De Clercq et al. 2013), but the species has not been detected in these countries to date. Considering the transparency of borders between Guinea, Liberia, and Sierra Leone, it can be assumed that *R. microplus* will be detected in these countries in a short time.

*Babesia bigemina* is common haemoparasite of cattle in Africa (Bock et al. 2004; Reye et al. 2012; Beckley et al. 2016; Kamani et al. 2017) and was detected in Nigeria (Ilemobade 1991), Ghana (Bell-Sakyi et al. 2004), and Guinea (De Meneghi et al. 2000). *Theileria parva* and *Theileria annulata* cause severe for domestic ruminants theileriosis transmitted by *Rhipicephalus appendiculatus* and by ticks which belong to the genus *Hyalomma* (Dolan 1989). Ticks of *Boophilus* species are not competent vectors of these two *Theileria* species, but engorged ticks might contain not only pathogens for which they are competent vectors but also other pathogens that circulate in the blood of the host at the time of tick feeding. For this reason, we screened collected ticks for the presence of *Theileria parva* and *Theileria annulata.* As a result, these pathogens were not detected in the collected ticks.

*Babesia bigemina* causes infections in livestock, and *R. microplus* is its competent vector. However, indigenous *Boophilus* species (*R. geigyi, R. annulatus*) are also able to transmit *B. bigemina* and had taken part in the pathogen circulation in West Africa before the invasion of *R. microplus* (Ilemobade 1991; Bell-Sakyi et al. 2004). According to Tomassone et al (2004), *B. bigemina* was found in 21.1% of blood smears taken from cattle in Guinea in 2000. No published research is available concerning tick-borne haemoparasites in ticks in Guinea, and we are not able to determine how the prevalence of the pathogens changed after the introduction of *R. microplus*.

*Anaplasma marginale* is a worldwide tick-borne disease of cattle that is characterized by progressive haemolytic anaemia, abortions, loss of condition, milk production, and death (Ristic 1981). In West Africa, *A. marginale* was detected in ticks and cattle in Senegal (Dahmani et al. 2019), Ivory Coast (Ehounoud et al. 2016), Benin (Adjou Moumouni et al. 2018a, b), and Nigeria (Ilemobade 1991; Reye et al. 2012; Lorusso et al. 2016). In Guinea, the prevalence of *A. marginale* in cattle in 2000 was 5.6% (Tomassone et al. 2004). Thus, the pathogen was presented in Guinea earlier, prior to the introduction of *R. microplus*. *Anaplasma platys* is primarily a pathogen of canines (Rymaszewska and Grenda 2008), but it was also found in cattle (Lorusso et al. 2016; Dahmani et al. 2019). Therefore, we hypothesize that the ticks obtained the pathogen from the blood of infected host animals.

We did not find *Ehrlichia ruminantium*, another important veterinary tick-borne pathogen, in our samples. Within the *Ehrlichia* genus, we found only uncultured genotypes in four ticks collected from the same cow. The sequences of 16S rRNA and groEl genes did not enable us to identify the pathogen at the species level. Further study of the molecular identification of this *Ehrlichia spp.* is needed.

## Conclusion

We reported the presence of *R. microplus* in Guinea and confirmed its further invasion in West African countries. The main route of species expansion is the transfer of cattle among countries without proper veterinary control. Most researchers who detected *R. microplus* in West Africa noted that the introduction of this species has negative consequences for livestock, primarily due to the transmission of vector-borne pathogens. We detected six pathogens in ticks, two of which, *B. bigemina* and *A. marginale*, cause severe diseases in cattle. However, both of these pathogens are common for West Africa and had been circulating in Guinea before the introduction of *R. microplus*. Nevertheless, the invasion of a new species of tick might change the composition of circulating pathogens and their prevalence. As a consequence, the introduction of new species might lead to considerable losses.

Clearly, it is not possible to prevent further distribution of *R. microplus* in West Africa within a climatically suitable range. Therefore, further studies of the pathogens circulating in these territories and the acaricidal resistance of ticks are warranted.

## Declarations

## Ethics approval

No ethics approval was required, as this study does not involve clinical trials or experimental procedures. The inspected cattle were slaughtered for human consumption. The slaughterhouse staff gave permission to collect ticks from animals. The Research Ethics Committee of the Central Research Institute of Epidemiology (Moscow, Russia) has confirmed that no ethics approval is required.

## Availability of data and material (data transparency)

The authors confirm that the data supporting the findings of this study are available within the article [.

